# Auricular vagus nerve stimulation facilitates contextual fear extinction via insular–mPFC pathway without affecting reconsolidation

**DOI:** 10.64898/2026.09.23.753025

**Authors:** Hiroshi Kuniishi, Eri Takeuchi, Okito Hashimoto, Mitsuhiko Yamada, Masayuki Sekiguchi, Hideo Matsuzaki

## Abstract

Cervical vagus nerve stimulation (VNS) suppresses conditioned fear in rodents and may serve as an adjunct to exposure-based therapy for anxiety disorders. Non-invasive auricular VNS also reduces conditioned fear in humans, although the underlying mechanisms remain unclear. Using a contextual fear conditioning paradigm, in which brief re-exposure induces reconsolidation and prolonged re-exposure promotes extinction learning in mice, we examined whether auricular VNS modulates these distinct memory processes. Auricular VNS during prolonged, but not brief, re-exposure reduced subsequent freezing, indicating facilitation of extinction learning without affecting reconsolidation. Stimulation restricted to the latter half of prolonged re-exposure, when extinction learning was underway, was sufficient to reduce subsequent fear responses. As a potential neural circuit mechanism, we focused on projections from the posterior insular cortex (pIC) to the medial prefrontal cortex (mPFC). The pIC receives interoceptive signals, including vagal input, whereas the mPFC plays a key role in fear regulation. Auricular VNS during extinction learning increased c-Fos expression in pIC neurons projecting to the infralimbic and dorsal peduncular cortex (IL/DP) within the mPFC. Optogenetic activation of the pIC– IL/DP pathway mimicked the extinction-facilitating effect of auricular VNS, whereas inhibition blocked it. These results demonstrate that auricular VNS promotes extinction learning of contextual fear memory by activating the pIC–IL/DP circuit, providing insight into how vagal afferent stimulation engages insular–prefrontal networks to modulate fear.

## Introduction

The brain and peripheral organs are engaged in complex bidirectional interactions that play a pivotal role in the regulation of their respective functions^1,2^. Among the principal mediators of this brain-body communication is the vagus nerve, which constitutes a major component of the parasympathetic nervous system. Through its afferent sensory fibres, the vagus nerve transmits signals reflecting the physiological status of peripheral organs to the central nervous system^3–7^. These afferent pathways not only help maintain homeostasis but also influence higher-order brain functions, including cognition, affective regulation, and emotional processing^8,9^. Consistent with this, several animal studies demonstrated the contribution of the vagus nerve to emotional regulation. For example, vagus nerve ablation alters innate anxiety-like behaviour and the extinction of conditioned fear in rodents^10^. Moreover, certain microbial strains and colonic inflammation modulate anxiety- and depression-like behaviours via vagus nerve-dependent mechanisms^11–13^. Together, these findings highlight the vagus nerve as a pivotal integrator of peripheral signals with central circuits underlying emotional regulation.

Clinically, vagus nerve stimulation (VNS) has been developed as a treatment for drug-resistant epilepsy^14^. Its therapeutic potential has been extended to psychiatric disorders, such as depression^15^. In rodent studies, cervical VNS also reduced conditioned fear responses^16–21^, suggesting its potential as an adjunct to exposure-based therapies for anxiety disorders, such as post-traumatic stress disorder (PTSD). More recently, attention has focused on the auricular branch of the vagus nerve, also known as Alderman’s or Arnold’s nerve, which innervates parts of the external auricle and projects to the nucleus tractus solitarius (NTS) in the brainstem^22,23^.

Based on this anatomical pathway, transcutaneous auricular vagus nerve stimulation (aVNS) has been developed as a non-invasive alternative to cervical VNS^24^, and several studies suggest that aVNS reduces conditioned fear responses also in humans^25–29^. However, the neural mechanisms underlying the effects of aVNS on fear regulation remain to be fully elucidated.

In the contextual fear conditioning paradigm, a brief re-exposure to a conditioned context induces fear reconsolidation, whereas prolonged re-exposures promote extinction learning^30–32^. Both reconsolidation blockade and extinction facilitation represent key strategies for reducing fear responses. By manipulating the duration of re-exposure, it is possible to dissociate whether an intervention on fear memory leads to attenuation or enhancement through the reconsolidation process or whether it facilitates extinction learning^33^. For example, a brief re-exposure session combined with midazolam interferes with the reconsolidation process and reduces the fear response. In this case, because brief re-exposure does not induce extinction learning, fear attenuation is considered to result from a disruption of the reconsolidation process rather than from extinction^34^. In contrast, the administration of an allosteric modulator of AMPA receptors, PEPA, combined with a short re-exposure session does not affect the fear response on the following day, whereas its administration with a prolonged re-exposure session reduces the subsequent fear response. These findings suggest that PEPA facilitates extinction learning rather than reconsolidation^33^. Moreover, D-cycloserine reduces fear responses when combined with a prolonged re-exposure session, whereas pairing D-cycloserine with an insufficiently short re-exposure can paradoxically increase subsequent fear responses, suggesting that D-cycloserine can facilitate extinction but also enhance reconsolidation when re-exposure is insufficient^33,35,36^. Thus, determining whether aVNS affects reconsolidation or extinction learning is essential for understanding how aVNS reduces conditioned fear.

To further understand how aVNS modulates conditioned fear, we next considered the neural circuits that may mediate its effects. Among the potential circuits, we focused on the projection from the posterior insular cortex (pIC), a region hypothesised to receive vagal afferent input, to the medial prefrontal cortex (mPFC), which plays a crucial role in fear regulation and extinction learning. The pIC receives interoceptive signals, including vagal input^8,37^, and human and animal studies demonstrated that cervical VNS can induce pIC activity^24,38^. In addition, previous studies demonstrated that fear extinction alters glucose metabolism, c-Fos expression, and real-time calcium responses in the pIC^39,40^ and that pharmacological inactivation of the pIC impairs conditioned safety discrimination^41^. These findings suggest an essential role of the pIC in fear extinction learning. Furthermore, the pIC sends dense projections to the infralimbic cortex (IL), a well-established region critical for fear extinction learning^42–44^, and to the dorsal peduncular cortex (DP), which has recently been implicated in fear encoding and extinction learning^45,46^, within the mPFC^47^. Notably, a human aVNS study reported that aVNS increases functional connectivity between the pIC and mPFC^48^. Together, these findings suggest the possibility that aVNS modulates fear extinction learning through activation of the pIC–IL/DP circuit.

Herein, we investigated whether aVNS suppresses fear responses by interfering with reconsolidation or by facilitating extinction learning in a contextual fear conditioning paradigm in mice, by varying the length of re-exposure sessions and timing of aVNS delivery. To examine the contribution of the pIC–IL/DP circuit to aVNS effects, we optogenetically manipulated transmission within this pathway during a fear extinction learning paradigm in mice.

## Results

### aVNS activates the NTS in behaving mice

To stimulate the auricular branch of the vagus nerve in behaving mice, we implanted T-shaped pin electrodes into the external ear and applied electrical stimulation. Based on previous studies, the electrodes were placed on the auricle such that stimulation of the helix (a region less innervated by the auricular branch of the vagus nerve) served as sham stimulation (Sham), whereas stimulation of the concha (a vagus-innervated region) was used as aVNS (Fig. 1a, b).

**Fig. 1.**
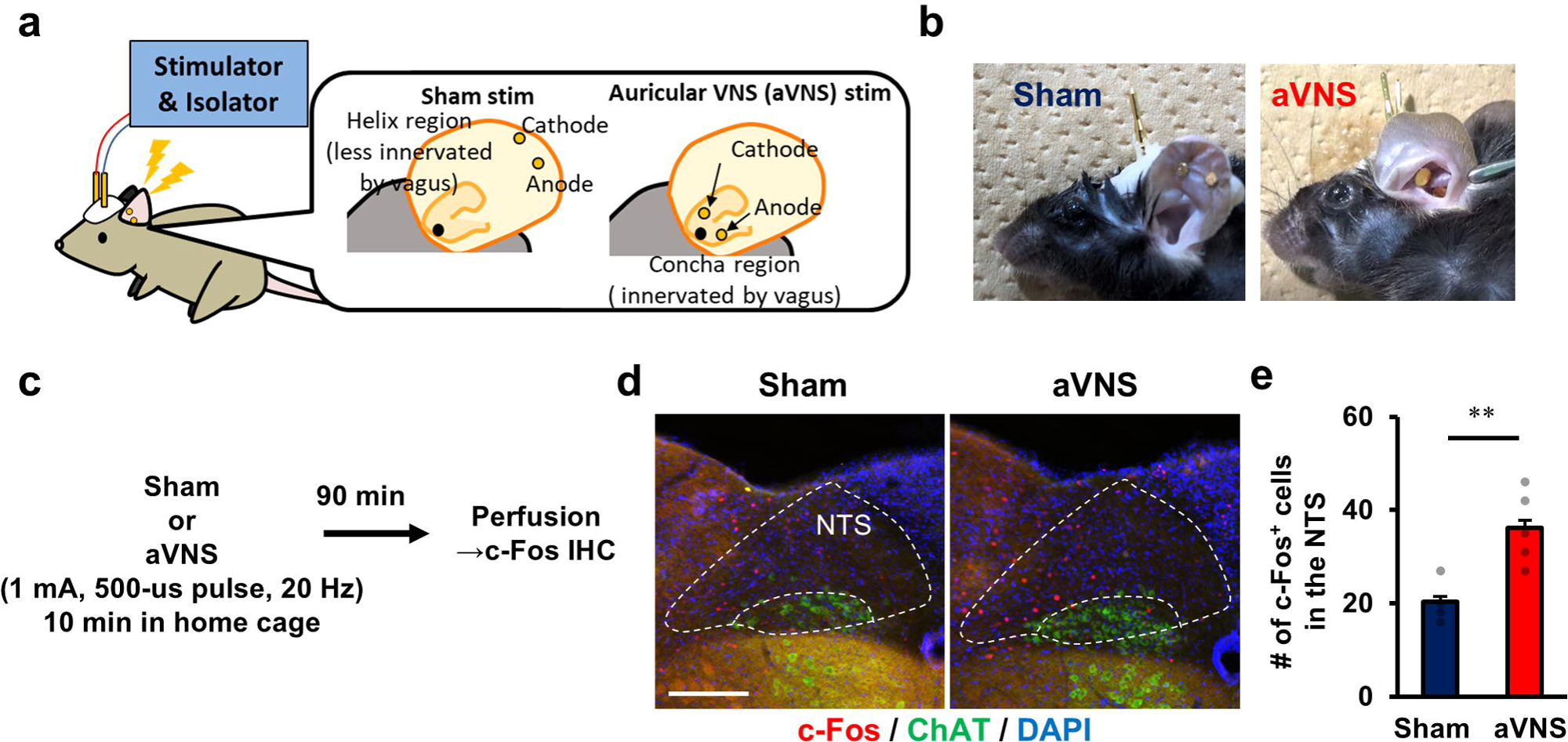
Auricular vagus nerve stimulation (aVNS) activates the nucleus tractus solitarius (NTS) in behaving mice. (**a**) Schematic illustration of electrode placement for auricular stimulation. For aVNS, electrodes were implanted in the concha region, which is innervated by branches of the vagus nerve. For sham stimulation (Sham), electrodes were implanted in the helix region, which is less innervated by vagal branches. Stimulation was delivered via an isolator and stimulator (1 mA, 500-μs pulse width, 20 Hz). (**b**) Representative photographs of mice with implanted electrodes. (**c**) Experimental timeline for c-Fos immunostaining following auricular stimulation. Mice received 10 minutes of stimulation in their home cages and were perfused 90 minutes later. IHC, immunohistochemistry. (**d**) Representative images of c-Fos and choline acetyltransferase (ChAT) immunostaining in the NTS. ChAT labels the dorsal motor nucleus of the vagus. Scale bar, 200 μm. DAPI, 4′,6-diamidino-2-phenylindole. (**e**) Quantification of c-Fos-positive cells in the NTS. The number of c-Fos-positive cells is significantly higher in the aVNS group than in the Sham group (unpaired t-test, t(7) = 3.564, p = 0.009). Group sizes: Sham (n = 4) and aVNS (n = 5). **p < 0.01

Afferent vagal fibres primarily project to the NTS in the medulla, and cervical VNS or aVNS has been reported to increase neural activity within the NTS^49–51^. To confirm the activation of vagal afferents, we applied electrical stimulation to the auricular region and performed c-Fos immunostaining in the NTS (Fig. 1c). Mice treated with aVNS showed a significant increase in the number of c-Fos-immunoreactive cells in the NTS, compared with Sham mice (Fig. 1d, e), consistent with the results of previous studies on aVNS^50,51^. These results suggest that our aVNS method activates vagal afferent input to the NTS in mice.

### aVNS facilitates extinction learning but does not interfere with the reconsolidation of contextual fear memory

To examine whether aVNS influences reconsolidation of fear memory or extinction learning, we applied this stimulation during either brief (3-minute) or prolonged (6-minute) re-exposure sessions and assessed freezing responses in subsequent retention tests. First, to examine the effect of aVNS on reconsolidation, we applied aVNS during brief re-exposure sessions (Fig. 2a). Freezing responses before and after conditioning on Day 1 did not differ between groups (Fig. 2b). When aVNS was delivered during the brief re-exposure session, a transient reduction in freezing was observed within this session compared with that in Sham controls (Fig. 2c).

**Fig. 2.**
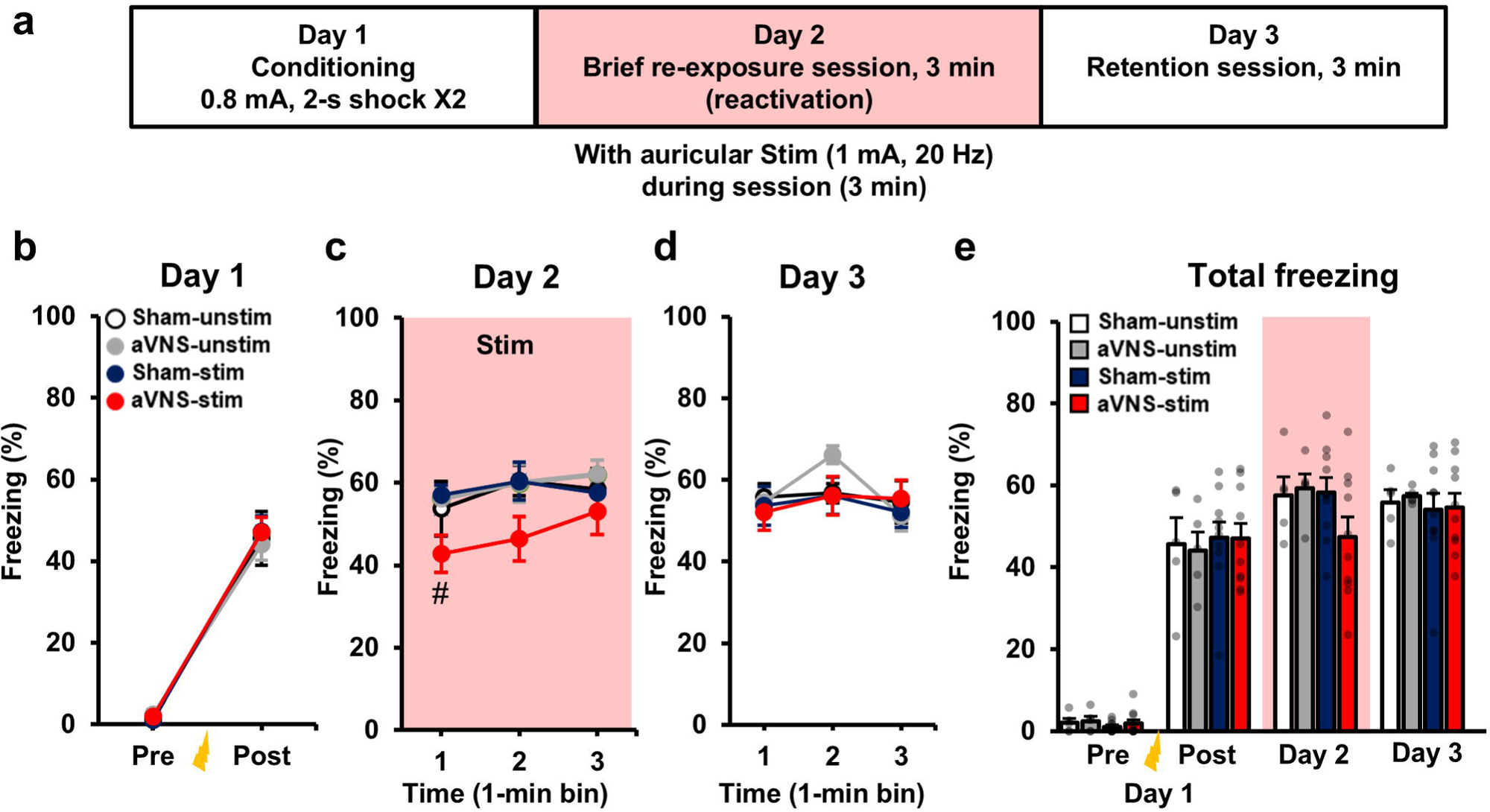
Auricular vagus nerve stimulation (aVNS) delivered during brief re-exposure does not disrupt the reconsolidation of contextual fear memory. (**a**) Experimental timeline. Mice underwent contextual fear conditioning with footshocks on day 1. On day 2, mice were re-exposed to the conditioned context for 3 minutes, without shocks, to reactivate the memory, and auricular stimulation was delivered during the re-exposure session (red shading). On day 3, mice were re-exposed to the same context for 3 minutes, without shocks or stimulation. (**b**) Freezing before and after footshocks on day 1. Repeated-measures analysis of variance (ANOVA) reveals no significant main effect of Treatment or Treatment × Bin interaction. (**c**) Freezing during the 3-minute re-exposure session on day 2, analysed in 1-minute bins. Repeated-measures ANOVA reveals a significant Treatment × Bin interaction (p = 0.012). Tukey’s honestly significant difference post hoc test shows a difference between aVNS-stim and Sham-stim (#p < 0.05). (**d**) Freezing during the day 3 retention test. Repeated-measures ANOVA reveals no significant main effect of Treatment or Treatment × Bin interaction. (**e**) Total freezing on day 1 (pre-and post-shock), 2, and 3. Repeated-measures ANOVA reveals no significant main effect of Treatment or Treatment × Session interaction. Group sizes: Sham-unstim (n = 5), aVNS-unstim (n = 5), Sham-stim (n = 10), and aVNS-stim (n = 10). Detailed statistical results are provided in **Supplementary Table 1**.

However, aVNS did not significantly decrease freezing in subsequent retention tests (Fig. 2d, e). Next, we applied aVNS during prolonged re-exposure sessions to examine its effects on extinction learning (Fig. 3a). Freezing responses before and after conditioning on Day 1 did not differ between groups (Fig. 3b). aVNS during the prolonged re-exposure significantly reduced freezing during extinction training and in retention tests on the following days, compared with Sham and non-stimulated controls (Fig. 3c–e). Together, these findings indicate that aVNS selectively enhances extinction learning without interfering with the reconsolidation of contextual fear memory.

**Fig. 3.**
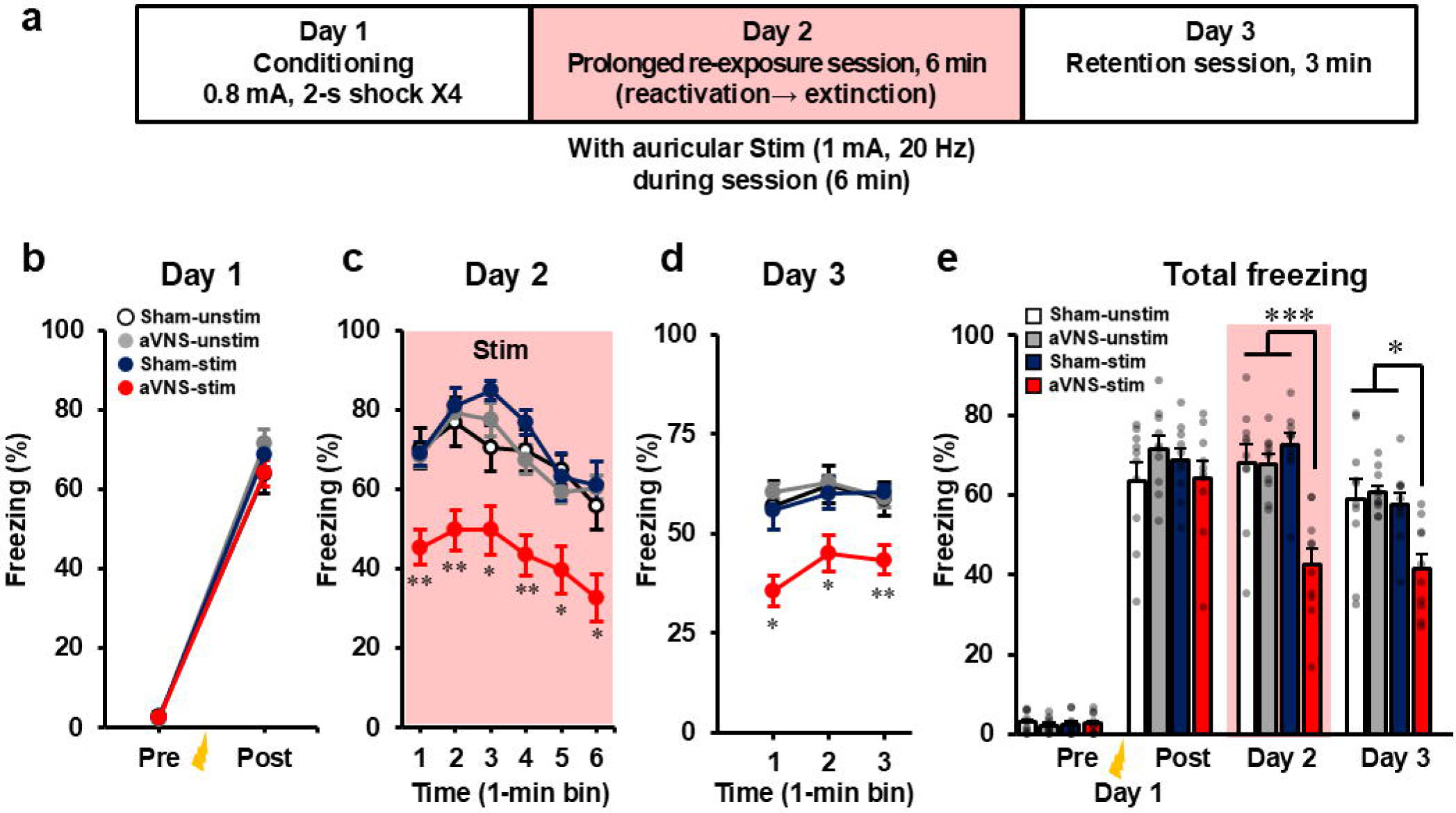
Auricular vagus nerve stimulation (aVNS) delivered during prolonged re-exposure facilitates extinction learning of contextual fear. (**a**) Experimental procedure. After contextual fear conditioning on day 1, mice were re-exposed to the conditioned context for 6 minutes on day 2 to induce extinction learning. Auricular stimulation was delivered during the re-exposure session (red shading). On day 3, mice were re-exposed to the same context, without shocks or stimulation. (**b**) Freezing before and after footshocks on day 1. Repeated-measures analysis of variance (ANOVA) reveals no significant main effect of Treatment or Treatment × Bin interaction. (**c**) Freezing during the day 2 re-exposure session (1-minute bins). Repeated-measures ANOVA reveals a significant main effect of Treatment (p < 0.001). Tukey’s honestly significant difference (HSD) post hoc test shows lower freezing rates in aVNS-stim mice than in the other groups (*p < 0.05, **p < 0.01). (**d**) Freezing during the day 3 extinction test. Repeated-measures ANOVA reveals a significant main effect of Treatment (p < 0.001). Tukey’s HSD test shows reduced freezing in aVNS-stim mice (*p < 0.05, **p < 0.01). (**e**) Total freezing across sessions. Repeated-measures ANOVA reveals a significant Treatment × Session interaction (p < 0.001). Tukey’s HSD test shows reduced freezing in aVNS-stim mice than in the other groups (*p < 0.05, ***p < 0.001). Group sizes: Sham-unstim (n = 10), aVNS-unstim (n = 10), Sham-stim (n = 10), and aVNS-stim (n = 10). Detailed statistical results are provided in **Supplementary Table 2**.

### aVNS in the latter half of prolonged re-exposure is sufficient to reduce fear responses

In our paradigm, mice show high freezing during the first 3 minutes of prolonged re-exposure, which gradually decreases during the latter 3 minutes, suggesting that extinction learning takes place during this phase^33^. Pharmacological interventions are difficult to apply within sessions, whereas aVNS allows precise timing of the stimulation. We hypothesised that, if aVNS suppresses fear by influencing extinction learning, then stimulation only during the latter half of the session would attenuate freezing in retention tests. Therefore, we applied aVNS during the latter or earlier half of a prolonged re-exposure session and assessed freezing responses in the retention sessions on the subsequent day (Fig. 4a, f). In the latter-half stimulation experiment, freezing responses before and after conditioning on Day 1 did not differ between groups (Fig. 4b). aVNS restricted to the latter half, a phase during which extinction learning is thought to occur, significantly decreased freezing during that phase and on the following test days, compared with Sham controls (Fig. 4c–e). In contrast, in the earlier-half stimulation experiment, freezing responses before and after conditioning on Day 1 did not differ between groups (Fig. 4g). Stimulation during the earlier half reduced freezing only transiently within the re-exposure session but did not affect retention (Fig. 4h–j). These findings suggest that the attenuation of contextual fear by aVNS critically depends on stimulation timing, with delivery during the late phase, when extinction learning is underway, being particularly important.

**Fig. 4.**
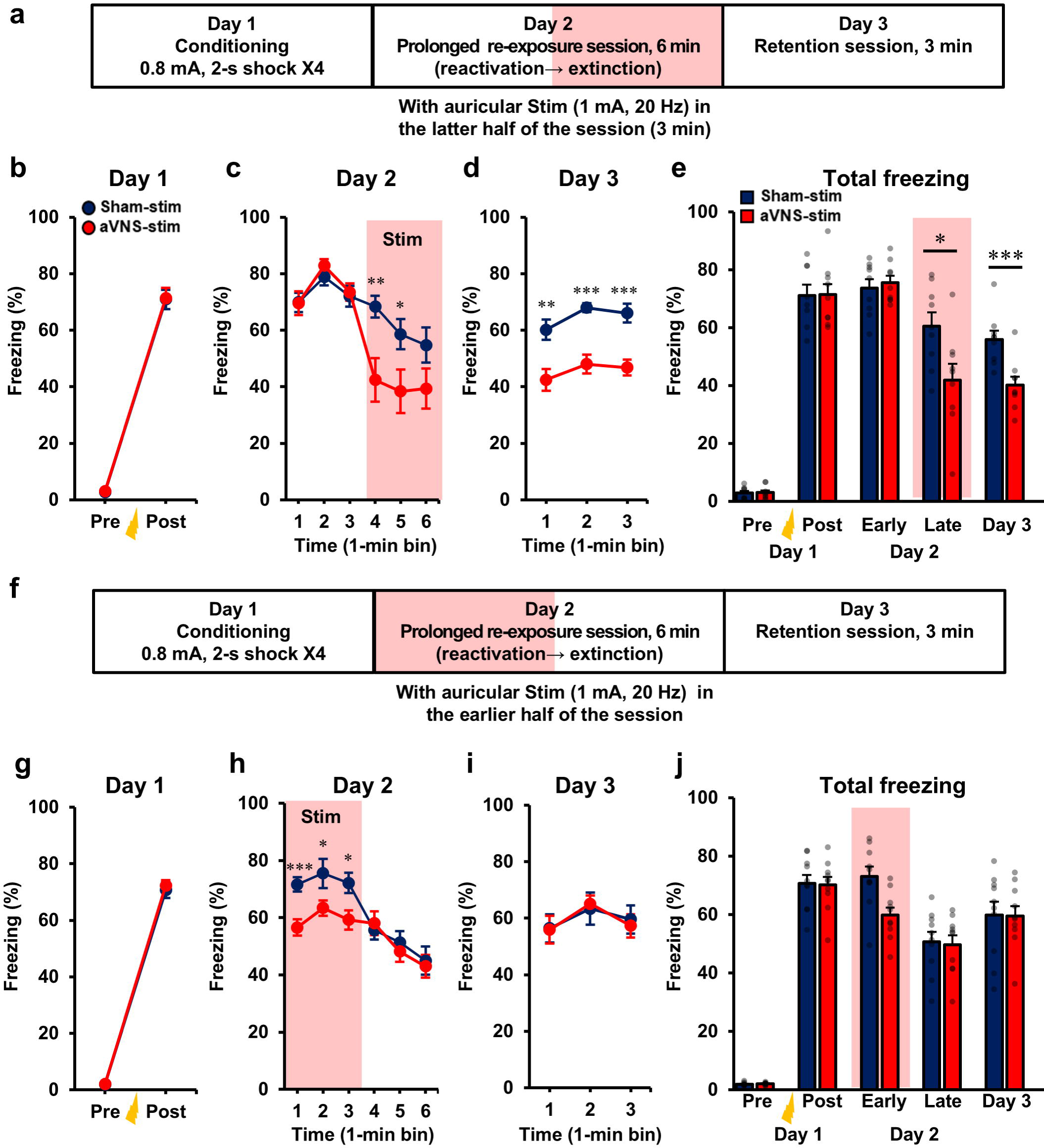
Auricular vagus nerve stimulation (aVNS) delivered during the latter half of a prolonged re-exposure session is sufficient to enhance extinction of contextual fear. (**a**) Experimental procedure (latter-half stimulation). Mice underwent contextual fear conditioning on day 1, followed by a 6-minute re-exposure session on day 2, without shocks. Auricular stimulation was delivered during the latter half of the session (red shading). On day 3, mice were re-exposed to the same context, without shocks or stimulation, to assess fear memory extinction. (**b**) Freezing before and after footshocks on day 1. Repeated-measures analysis of variance (ANOVA) shows no main effect of Treatment or Treatment × Bin interaction. (**c**) Freezing during the day 2 re-exposure session, analysed in 1-minute bins. Repeated-measures ANOVA reveals a significant Treatment × Bin interaction (p < 0.001). Post hoc unpaired t-tests demonstrate significant differences at 4 and 5 minutes (*p < 0.05, **p < 0.01). (**d**) Freezing during the day 3 test, analysed in 1-minute bins. Repeated-measures ANOVA shows a significant main effect of Treatment (p < 0.001). Post hoc comparisons were performed only when a significant interaction or main effect was detected (**p < 0.01, ***p < 0.001). (**e**) Total freezing across sessions. Repeated-measures ANOVA reveals a significant Treatment × Session interaction. Post hoc comparisons were performed only when a significant interaction or main effect was detected (*p < 0.05, ***p < 0.001). Group sizes: Sham-stim (n = 9) and aVNS-stim (n = 9). (**f**) Experimental procedure (earlier-half stimulation). (**g–j**) With earlier-half stimulation, freezing during the day 2 re-exposure session (**h**) shows a significant Treatment × Bin interaction, with significant early time bins identified by post hoc unpaired t-tests (*p < 0.05, ***p < 0.001), whereas day 3 freezing (**i**) and total freezing (**j**) show no significant effects of Treatment (p > 0.05). Group sizes: Sham-stim (n = 10) and aVNS-stim (n = 10). Detailed statistical results are provided in **Supplementary Table 3**.

### aVNS activates the pIC–IL/DP pathway

Visceral sensory information transmitted via the vagus nerve is thought to be conveyed to the NTS and then relayed through the parabrachial nucleus and the thalamus to the pIC^8,9^. Moreover, the pIC sends specific projections to the IL and DP in the mPFC^47^. The IL is a critical brain region for fear extinction learning, and the DP has recently been implicated in fear encoding and fear extinction. Hence, we hypothesised that aVNS facilitates fear extinction learning through enhanced excitability of the pIC–IL/DP circuit. To confirm pIC–IL/DP projections, we injected the anterograde tracer Fluoro-Ruby into the pIC. Anterogradely labelled axon terminals were preferentially observed in the IL and DP but not the prelimbic cortex, in the mPFC (Fig. S1a–d). Next, to examine whether aVNS applied during fear extinction learning activates pIC neurons that project to the IL or DP, we injected a retrograde green fluorescent protein (GFP)-expressing adeno-associated virus (AAV) vector into the IL/DP and assessed c-Fos expression in the retrogradely labelled pIC neurons (Fig. 5a–c). After several weeks to allow for GFP expression, mice were perfused following the administration of aVNS during the latter half of the prolonged re-exposure session, and the expression of GFP and c-Fos was examined (Fig. 5d). The total numbers of GFP- and c-Fos-expressing cells in the pIC did not significantly differ between aVNS and Sham treatments (Fig. 5e–g). Notably, the proportion of GFP-positive cells co-expressing c-Fos was significantly higher in the aVNS group than in the Sham group (Fig. 5e, h). These results suggest that aVNS during extinction learning preferentially activates pIC–IL/DP projecting neurons.

**Fig. 5.**
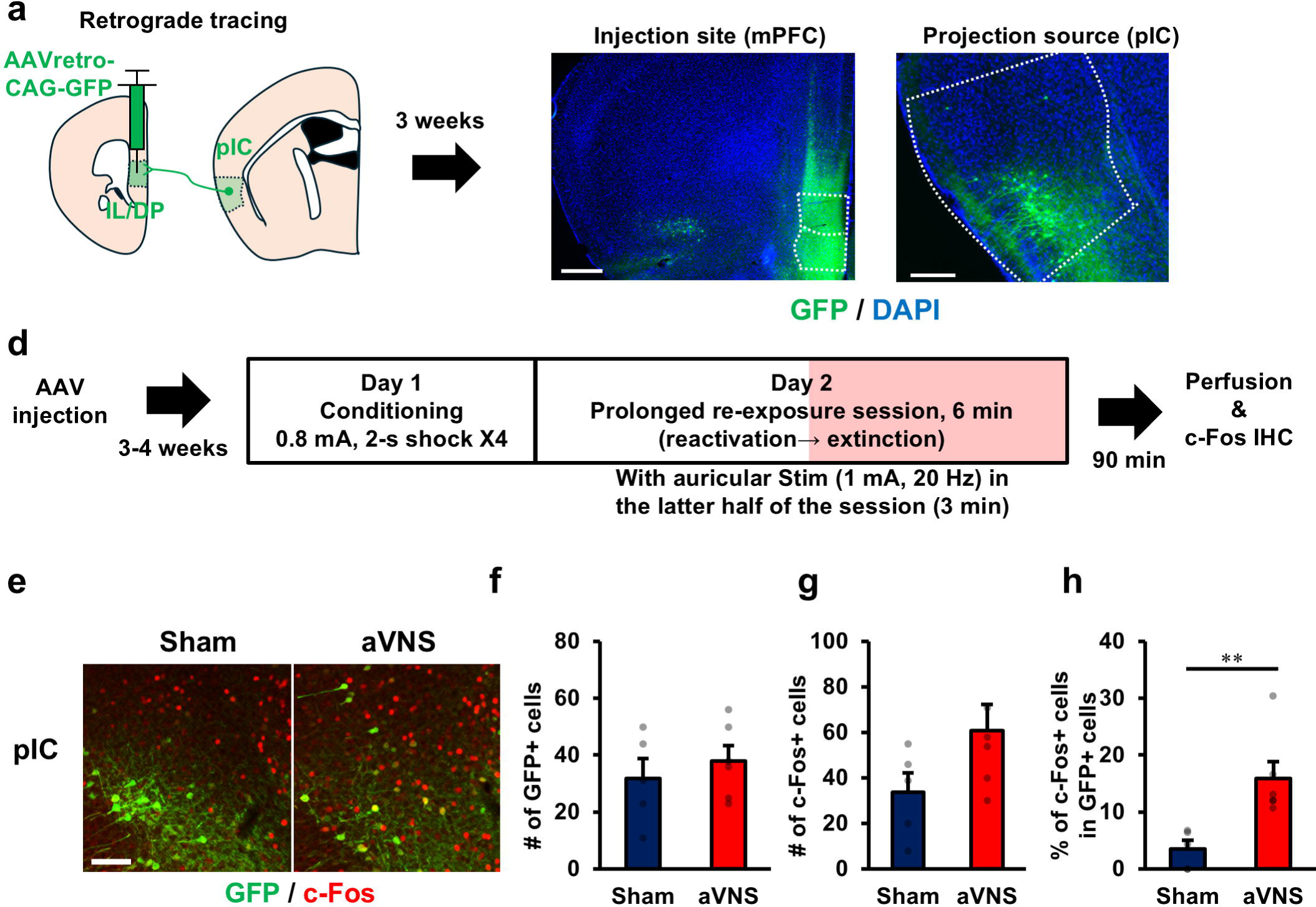
Auricular vagus nerve stimulation (aVNS) delivered during the latter half of a prolonged re-exposure session activates neurons projecting from the posterior insular cortex (pIC) to the infralimbic and dorsal peduncular cortex (IL/DP). (**a**) Schematic describing the retrograde tracing of pIC→IL/DP projection neurons. (**b**) Representative image of the retrograde adeno-associated virus (AAV) injection site in the IL/DP region of the medial prefrontal cortex (mPFC). Scale bar, 500 μm. (**c**) Representative image of green fluorescent protein (GFP)-labelled IL/DP-projecting neurons in the pIC. Scale bar, 250 μm. DAPI, 4′,6-diamidino-2-phenylindole. (**d**) Experimental procedure for c-Fos analysis following auricular stimulation. IHC, immunohistochemistry. (**e**) Representative images of GFP and c-Fos immunostaining in the pIC. (**f**) The number of GFP-labelled projection neurons in the pIC does not differ between groups (unpaired t-test, t(9) = 0.713, p = 0.494). (**g**) The number of c-Fos-positive neurons in the pIC does not differ between groups (unpaired t-test, t(9) = 1.802, p = 0.105). (**h**) Percentage of c-Fos-positive neurons among GFP-labelled neurons is increased in aVNS mice compared with Sham mice (unpaired t-test, t(9) = 3.310, p = 0.008). Group sizes: Sham (n = 5) and aVNS (n = 6). **p < 0.01.

### Optogenetic activation of the pIC–IL/DP pathway enhances fear extinction learning

Next, we examined whether the activation of the pIC–IL/DP pathway during extinction learning enhances contextual fear extinction learning, similar to aVNS. We injected an AAV vector expressing channelrhodopsin-2 (ChR2) under the control of the CaMKII promoter into the pIC of mice and implanted an optical fibre above the IL/DP to deliver blue light-emitting diode (LED) light (Fig. 6a, b and Fig. S2a, b). Using these mice, we activated the projection pathway from the pIC to the IL/DP by delivering optical stimulation during the latter half of the prolonged re-exposure session (Fig. 6c). In this experiment, freezing responses before and after conditioning on Day 1 did not differ between groups (Fig. 6d). Optogenetic stimulation during the latter half of the prolonged re-exposure session significantly decreased freezing during that phase and on the following test days, compared with enhanced yellow fluorescent protein (EYFP)-expressing controls (Fig. 6e–g). These results indicate that activation of the pIC–IL/DP pathway, similar to aVNS, facilitates fear extinction learning in mice.

**Fig. 6.**
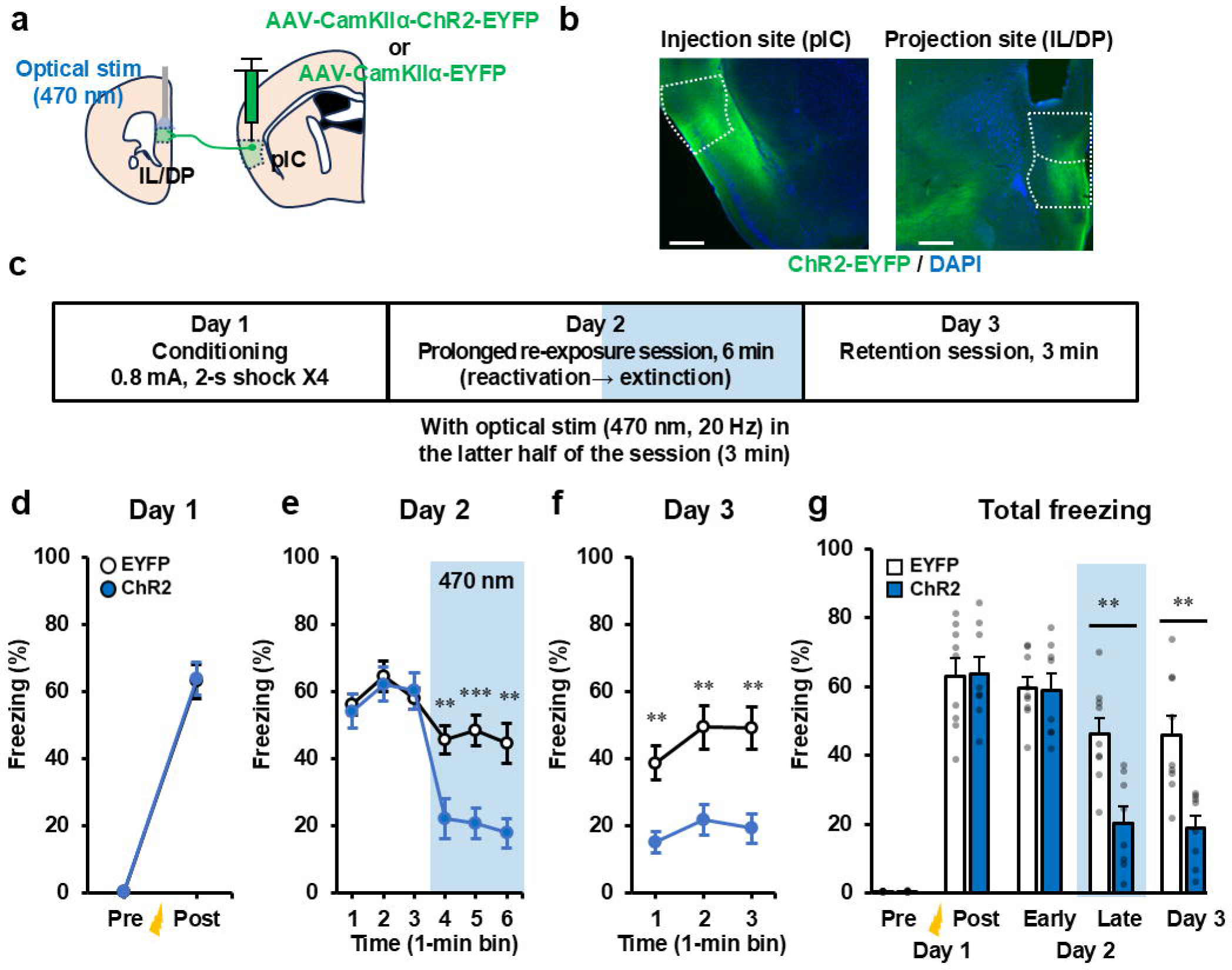
Optogenetic activation of the posterior insular cortex (pIC)→ infralimbic/dorsal peduncular cortex (IL/DP) pathway during the latter half of a prolonged re-exposure session is sufficient to enhance extinction of contextual fear. **(a)** Schematic of optogenetic activation experiment. AAV-CamKIIα-ChR2-EYFP or control AAV-CamKIIα-EYFP was injected into the pIC. Optical fibres were implanted above the IL/DP to activate pIC→IL/DP axon terminals (470 nm). (**b**) Representative images show viral injection site in the pIC and EYFP-labelled projections in the IL/DP. Scale bar, 500 μm. (**c**) Experimental procedure. After contextual fear conditioning with footshocks on day 1, mice were re-exposed to the conditioned context for 6 minutes on day 2, without shocks, to induce extinction learning. Optical stimulation (470 nm) of pIC axon terminals in the IL/DP was delivered during the latter half of the re-exposure session (blue shading). On day 3, mice were re-exposed to the same context for 3 minutes, without shocks or stimulation, to assess fear memory extinction. (**d**) Freezing before and after footshocks on day 1. Repeated-measures analysis of variance (ANOVA) shows no main effect of Treatment or Treatment × Bin interaction. (**e**) Freezing during the 6-minute re-exposure session on day 2, analysed in 1-minute bins. Repeated-measures ANOVA shows significant main effect of Treatment and Treatment × Bin interaction. Post hoc unpaired t-tests show reduced freezing in channelrhodopsin-2 (ChR2)-expressing mice at 4–6 minutes (**p < 0.01, ***p < 0.001). (**f**) Freezing during the extinction test on day 3, analysed in 1-minute bins. Repeated-measures ANOVA shows a significant main effect of Treatment. Post hoc unpaired t-tests show reduced freezing in ChR2-expressing mice (**p < 0.01). (**g**) Total freezing across sessions. Repeated-measures ANOVA revealed a significant Treatment × Session interaction. Post hoc unpaired t-tests show reduced freezing in ChR2-expressing mice during late extinction and on day 3 (**p < 0.01). Group sizes: EYFP (n = 9) and ChR2 (n = 8). Detailed statistical results are provided in **Supplementary Table 4**.

### Optogenetic inhibition of the pIC–IL/DP pathway during aVNS prevents its facilitating effect on fear extinction

To further investigate the contributions of the pIC–IL/DP pathway to the effects of aVNS, we optogenetically inhibited the pIC–IL/DP pathway during aVNS application in the extinction learning phase. For the inhibition of the pIC–IL/DP pathway, we injected an AAV vector expressing halorhodopsin (NpHR) under the control of the CaMKII promoter into the pIC of mice and implanted an optical fibre above the IL/DP to deliver yellow LED light (Fig. 7a, b and Fig. S3a, b). Using these mice, we inhibited the pIC–IL/DP pathway by delivering optical stimulation during the latter half of the prolonged re-exposure session with aVNS (Fig. 7c). In this experiment, freezing responses before and after conditioning on Day 1 did not differ between groups (Fig. 7d). During the latter half of the prolonged re-exposure session, aVNS significantly decreased freezing responses in EYFP-expressing mice, and this reduction persisted in the retention test on the following day. In contrast, these effects of aVNS were abolished in NpHR-expressing mice, in which the pIC–IL/DP pathway was inhibited during aVNS (Fig. 7e– g). Taken together, these findings indicate that activation of the pIC–IL/DP pathway is essential for the fear extinction-facilitating effects of aVNS.

**Fig. 7.**
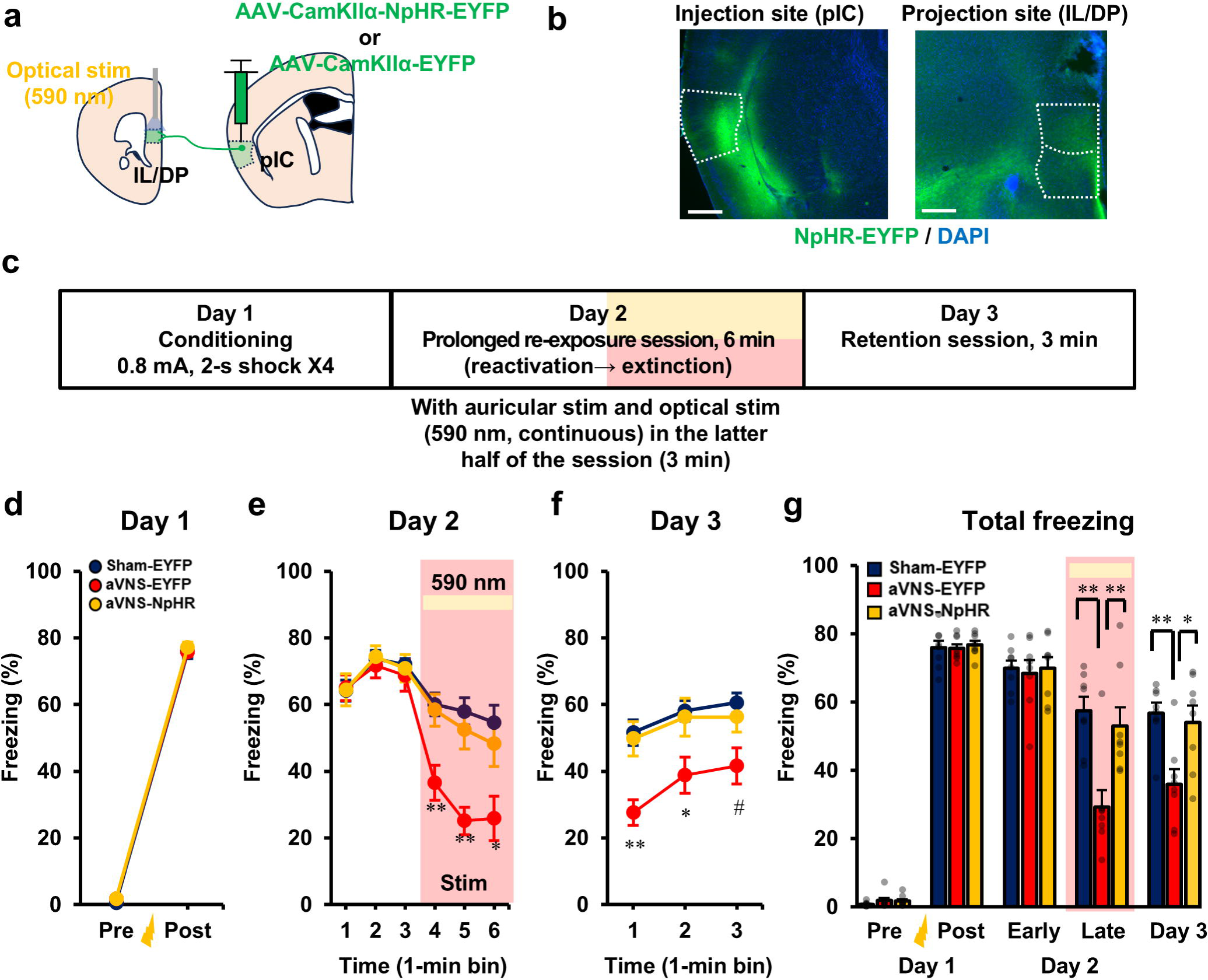
Optogenetic inhibition of the posterior insular cortex (pIC)=→ infralimbic/dorsal peduncular cortex (IL/DP) pathway blocks auricular vagus nerve stimulation (aVNS)-induced enhancement of contextual fear extinction. (**a**) Schematic of optogenetic inhibition experiment. AAV-CamKIIα-NpHR-EYFP or control AAV-CamKIIα-EYFP was injected into the pIC. Optical fibres were implanted above the IL/DP to inhibit pIC→IL/DP axon terminals (590 nm). (**b**) Representative images show viral injection site in the pIC and EYFP-labelled projections in the IL/DP. Scale bar, 500 μm. (**c**) Experimental timeline. Mice underwent contextual fear conditioning on day 1. On day 2, mice were re-exposed to the conditioned context for 6 minutes, without shocks, to induce extinction learning. aVNS was delivered during the latter half of the session, with optical inhibition of pIC→IL/DP terminals. On day 3, mice were re-exposed to the same context, without shocks or stimulation, to assess fear memory extinction. (d) Freezing before and after footshocks on day 1. Repeated-measures analysis of variance (ANOVA) shows no main effect of Treatment or Treatment × Bin interaction. (**e**) Freezing during the 6-minute re-exposure session on day 2, analysed in 1-minute bins. Repeated-measures ANOVA shows significant main effect of Treatment and Treatment × Bin interaction. Tukey’s honestly significant difference (HSD) post hoc test shows reduced freezing in aVNS-EYFP mice compared with the other groups (*p < 0.05, **p < 0.01). (**f**) Freezing during the extinction test on day 3, analysed in 1-minute bins. Repeated-measures ANOVA shows a significant main effect of Treatment. Tukey’s HSD post hoc test shows reduced freezing in aVNS-EYFP mice compared with the other groups (*p < 0.05, **p < 0.01) and a difference between Sham-EYFP and aVNS-EYFP mice (#p < 0.05). (**g**) Total freezing across sessions. Repeated-measures ANOVA reveals a significant Treatment × Session interaction. Tukey’s HSD post hoc test shows reduced freezing in aVNS-EYFP mice during late extinction and the day 3 test, compared with the other groups (*p < 0.05, **p < 0.01). Group sizes: Sham-EYFP (n = 8), aVNS-EYFP (n = 8), and aVNS-NpHR (n = 8). Detailed statistical results are provided in **Supplementary Table 5**.

## Discussion

In our model, the application of aVNS during short re-exposure sessions that induce reconsolidation did not affect fear responses on the following day. In contrast, stimulation delivered throughout prolonged re-exposure sessions that promote extinction learning reduced fear responses in subsequent tests. Notably, restricting stimulation to the latter half of prolonged re-exposure, when extinction learning is presumed to be most active, was sufficient to produce this effect. These findings indicate that aVNS attenuates subsequent fear responses not by disrupting reconsolidation but by enhancing extinction learning. Our anatomical and optogenetic analyses further revealed that this extinction-facilitating effect involves the activation of pIC neurons projecting to the IL/DP. Together, these findings identify the pIC–IL/DP pathway as a critical circuit underlying the extinction-enhancing effects of aVNS and establish a time- and circuit-specific mechanism through which aVNS modulates fear memory.

In addition to aVNS, several pharmacological interventions facilitate fear extinction under conditions of sufficient re-exposure. For example, D-cycloserine and certain histone deacetylase inhibitors can enhance extinction learning. However, when re-exposure is insufficient, these compounds may strengthen reconsolidation and paradoxically exacerbate subsequent fear responses^35,36,52^. Such paradoxical effects raise concerns for the clinical application of this intervention because suboptimal exposure conditions may lead to adverse outcomes. Our data indicate that aVNS does not interfere with reconsolidation processes and instead facilitates extinction without exacerbating subsequent fear expression. This feature highlights a potential advantage of aVNS as a neuromodulatory approach that can be safely combined with exposure-based therapies, thereby enhancing extinction learning while minimising the risk of inadvertently strengthening maladaptive fear memories.

A key advantage of electrical neuromodulation over pharmacological approaches is its precise temporal control. Leveraging this feature, we show that restricting aVNS to the late phase of prolonged re-exposure is sufficient to facilitate extinction learning. In contrast, stimulation confined to the early phase produces no lasting effect. These findings indicate that aligning VNS with extinction-related neural plasticity and associated cognitive and physiological processes is critical for optimal efficacy.

Building on the importance of temporal alignment, closed-loop stimulation systems have been proposed, in which neural states are monitored in real time, for example using electroencephalography, and stimulation is delivered only when needed^53^. Identifying biomarkers for extinction-active phases may enable the delivery of aVNS within optimal temporal windows. Although aVNS is generally considered safe, mild adverse effects, such as transient discomfort, skin irritation, headache, and dizziness, have been reported^54^. Implementing a closed-loop system that delivers stimulation only at the most appropriate timing might further minimise these unwanted effects while enhancing patient comfort and compliance. Moreover, such a timing-precise state-dependent stimulation strategy may maximise therapeutic specificity and efficacy, paving the way for personalised and adaptive neuromodulation approaches in the treatment of fear- and anxiety-related disorders.

In the present study, aVNS transiently reduced freezing when delivered during re-exposure sessions, including brief re-exposure and the early or late phases of prolonged sessions. However, only stimulation delivered under extinction-permissive conditions, either throughout prolonged re-exposure or during its late phase, produced persistent attenuation of fear responses on the following day. These findings indicate that the immediate within-session suppression of freezing and the enduring reduction observed on subsequent days can be dissociated. This dissociation raises the possibility that aVNS engages partially separable processes, including an acute, state-dependent modulation of fear expression and a learning-related mechanism that facilitates extinction when stimulation coincides with extinction-associated plasticity.

The insular cortex is a core region for interoception, integrating visceral signals into representations of bodily states that shape affective and cognitive processes^55^. In particular, the pIC is thought to receive vagal afferent input via the NTS, parabrachial nucleus, and thalamus and to convey this information to higher-order brain regions involved in emotional regulation, such as the mPFC. The IL, a subdivision of the mPFC, has long been recognised as a critical structure for fear extinction learning^42,44^, whereas the DP, another mPFC subdivision, has more recently been implicated in the encoding of fear and the regulation of extinction learning^45,46^. In the present study, we demonstrated the importance of the pIC–IL/DP pathway in the facilitation of fear extinction induced by aVNS. Our findings extend previous knowledge by highlighting the contribution of the pIC from the perspective of interoceptive processing within anatomically defined neural circuits. A previous study demonstrated that projections from the pIC to the central amygdala contribute to fear acquisition, whereas projections to the nucleus accumbens have been implicated in fear extinction^56^. Herein, we further demonstrated that projections from the pIC to the mPFC are required for the extinction-facilitating effects of aVNS. These findings advance our understanding of pathway-specific mechanisms by which circuits downstream of the pIC differentially regulate fear acquisition and extinction.

The antiepileptic effects of VNS have been attributed to activation of the locus coeruleus– noradrenergic (LC–NA) system^57^, which also contributes to fear consolidation and extinction^58–60^. In rodent studies, cervical VNS increases central noradrenaline release^61^, and optogenetic inhibition of LC–NA neurons abolishes the extinction-facilitating effects of cervical VNS^62^, suggesting the involvement of the LC–NA system in the fear-regulating effects of VNS. In the present study, we identify the pIC–mPFC pathway, from an interoceptive perspective, as an additional circuit contributing to the facilitation of fear extinction by VNS. The relationship between this pathway and the LC–NA system remains unclear. However, adrenergic receptors are expressed in the insular cortex and the mPFC, and adrenergic signalling in these regions has been implicated in their functional regulation^63–65^. Thus, LC-derived NA modulation might influence fear extinction by regulating the activity of the pIC–mPFC circuit. Further studies are required to clarify how these systems interact.

The present study has some limitations. We applied aVNS using a single stimulation parameter determined based on prior studies^66,67^. Studies of cervical VNS have reported that the extinction-facilitating effects of stimulation follow a U-shaped function of intensity^68^, with certain levels being beneficial, whereas others may even exert detrimental effects. The efficacy of VNS also varies depending on stimulation frequency^69^. In human studies of VNS, both positive and null effects on fear extinction have been reported, which may be attributable to differences in stimulation parameters^25,27,28,70^. Therefore, it will be important to clarify how stimulation parameters, such as intensity, frequency, and laterality, influence fear extinction and potential unwanted effects, to define the effective range of stimulation. Establishing these optimal stimulation conditions will be critical not only for maximising therapeutic efficacy but also for minimising adverse outcomes. Such knowledge will be essential for the safe and effective clinical application of aVNS and for the development of individualised neuromodulatory strategies for treating fear-related disorders, such as PTSD and anxiety disorders.

## Methods

### Animals

Male C57BL/6J mice (8–12 weeks old at the time of behavioural testing and histological analyses) were purchased from CLEA Japan (Tokyo, Japan). Mice were housed in groups of four per plastic cage under controlled temperature (22 ± 1°C) and lighting (12-hour light/dark cycle, lights on at 08:00 a.m.) conditions, with *ad libitum* access to food and water. All experiments were conducted in strict accordance with the regulations of the University of Fukui and the National Institute of Neuroscience (Japan) for animal experimentation (approval numbers: 2020025, 2021037, R04090, R05074, R06066, and R07058).

### Surgery for aVNS

Mice were anaesthetised by intraperitoneal injection of medetomidine (0.75 mg/kg), midazolam (4 mg/kg), and butorphanol (5 mg/kg). After a midline scalp incision, two T-shaped brass pins (Daiso, Higashi-Hiroshima, Japan) were inserted for aVNS through the concha (innervated by the vagus nerve) or for Sham stimulation through the helix (less innervated by the vagus nerve) of the left external ear (Fig. 1a, b) and fixed in place using earring backings. Electrode locations were determined based on the results of previous rodent studies^66,67,71^. The pins were soldered to insulated copper wires placed subcutaneously. The opposite ends of the wires were soldered to male miniature connectors (A-M Systems, Sequim, WA, USA), which were then fixed to the skull with dental cement (Fig. 1a). During soldering, the external ear was cooled with ice to prevent thermal damage. After surgery, mice were intraperitoneally injected with atipamezole (0.75 mg/kg), housed individually, and allowed to recover for 3–5 days before behavioural testing or histological analysis.

### aVNS in freely behaving mice

For aVNS, mice were gently restrained, and female metallic connectors attached to an electrical stimulator and isolator (Nihon Kohden, Tokyo, Japan) were connected to the male pins fixed to the skull. Electrical stimulation consisted of 20-Hz uniphasic square pulses (1 mA, 500 µs pulse width, positive polarity) delivered under the control of the stimulator and isolator. The stimulation parameters were based on previous studies^66,67^.

### Immunostaining

Mice were transcardially perfused with phosphate-buffered saline (PBS), and the brains were removed and post-fixed. Coronal brain sections (50-µm thick) were cut using a vibratome. Free-floating sections were incubated in PBS containing 5% normal donkey serum (Vector Laboratories, Burlingame, CA, USA) and 0.3% Triton X-100 for 1 hour at room temperature and then incubated overnight at room temperature with primary antibodies against c-Fos (1:2,000; #2250, Cell Signaling Technology, Danvers, MA, USA), choline acetyltransferase (1:200; AB144P, Millipore, Burlington, MA, USA), and GFP (1:1,000; GF090R, Nacalai Tesque, Kyoto, Japan). After rinsing in PBS, sections were then incubated for 2 hours at room temperature with Alexa Fluor 488-(1:500; A-21208 or A-11055, Invitrogen, Carlsbad, CA, USA) or 594-conjugated secondary antibodies (1:500; A-21207, Invitrogen) and 4′,6-diamidino-2-phenylindole.

### Contextual fear conditioning and extinction training

Contextual fear conditioning was performed as previously described^33^. On day 1 (conditioning), mice were placed in the conditioning chamber and allowed to explore for 148 seconds, followed by 2 footshocks (0.8 mA, 2 s; mean inter-shock interval: 28 s) for reconsolidation experiments or 4 footshocks for extinction experiments. For reconsolidation experiments, we used the milder conditioning protocol (two shocks) to avoid ceiling effects and to facilitate detection of bidirectional effects on reconsolidation (enhancement and attenuation). Mice were removed 60 seconds after the final shock and returned to their home cages. On day 2 (extinction training), mice were re-exposed to the chamber for 3 (for reconsolidation) or 6 (for extinction) minutes without footshocks, with either aVNS or Sham stimulation. On day 3 (retention test), mice were re-exposed to the chamber for 3 minutes without shocks or stimulation.

### Stereotaxic surgery for tracer studies, viral injection, and optic cannula implantation

Stereotaxic surgery was performed as previously described^72^. Mice were anaesthetised with medetomidine, midazolam, and butorphanol via intraperitoneal injection and secured in a stereotaxic apparatus (Narishige, Tokyo, Japan). Eyes were protected from drying with ophthalmic ointment (Tarivid; Santen Pharmaceutical, Osaka, Japan). Small craniotomies were made above the IL/DP (2.0 mm anterior to bregma, 0.25 mm lateral) or pIC (0.5 mm posterior to bregma, 3.7 mm lateral). Viral vectors (AAVrg-CAG-GFP, AAV5-CaMKIIa-EYFP, AAV5-CaMKIIa-ChR2-EYFP, or AAV5-CaMKIIa-eNpHR3.0-EYFP; University of North Carolina Vector Core and Addgene) or anterograde neuronal tracer (Fluoro-Ruby; Thermo Fisher Scientific, Waltham, MA, USA, 10% in saline) were pressure-injected (300 nL, 100 nL/min) at depths of 1.8 mm (pIC) or 2.0 mm (IL/DP) using a 10-µL Hamilton syringe driven by an infusion pump (UMP-3; World Precision Instruments, FL, USA). The needle was left in place for 5 minutes before slow retraction. After surgery, mice received an intraperitoneal injection of atipamezole. After 3–6 weeks, mice were used for behavioural experiments. For optogenetic manipulation, dual-LED optic cannulae (TeleLCD-B-500-0.8 or TeleLCD-Y-5-500-0.8; BRC Nihon Bioresearch, Hashima, Japan) were implanted bilaterally in the IL/DP (2.0 mm anterior to bregma, 0.4 mm lateral, −1.7 mm from the dura) and fixed to the skull with dental cement. After surgery, mice were intraperitoneally injected with atipamezole. After behavioural testing, mice were intracardially perfused, the brains were removed, and the locations of injections and cannulae were histologically verified.

### Optogenetic stimulation in behaving mice

To manipulate the pIC–IL/DP pathway, ChR2 or NpHR was expressed under the CaMKII promoter in the pIC. Control mice expressed EYFP. Optogenetic stimulation was performed wirelessly (Teleopto; BRC Nihon Bioresearch) through LED optic cannulae implanted above the IL/DP. Before testing, an infrared receiver (TeleR-2-P) was connected to the LED cannula. Blue light (470 nm, 10–16 mW, 10-millisecond pulses at 20 Hz) for ChR2 or yellow light (590 nm, 7– 14 mW, continuous) for NpHR was delivered under the control of a remote controller (TeleRemocon) and stimulator (SEN-7203; Nihon Kohden). In the contextual fear conditioning paradigm, optical stimulation was applied during the latter 3 minutes of the 6-minute re-exposure session on day 2.

### Statistical analysis

Data are presented as mean ± standard error of the mean. All analyses were performed using EZR. Comparisons between the two groups were performed using a two-tailed unpaired Student’s t-test. For multiple comparisons, one-way analysis of variance (ANOVA) followed by Tukey’s honestly significant difference (HSD) test was used. For repeated-measures data with two factors, two-way repeated-measures ANOVA was performed. When a significant main effect or interaction was detected, post hoc pairwise comparisons were performed using the unpaired two-tailed Student’s t-test or Tukey’s HSD test. A p value < 0.05 was considered significant. The exact statistical values are provided in Supplementary Tables 1–5.

## Data availability

Source data are provided with this paper. Additional data that support the findings of this study are available from the corresponding author upon reasonable request.

## Acknowledgments

We would like to thank Ms. Natsuki Miyakoshi and Ms. Hiromi Fujita for assistance with the animal care, and Mr. Riku Adachi, Ms. Yasuko Nakamura, and Ms. Ami Kakimoto for assistance with the experiments.

## Author contributions

H.K., O.H., and M.S. contributed to the conceptualization and design of the experiments. H.K. and E.T. contributed to performance of the experiments and analysis of the data. M.S., M.Y., and H.M. contributed to the development of experimental equipment. H.K. contributed to writing of the manuscript. All authors discussed the results and their interpretation and reviewed the manuscript.

## Declaration of generative AI and AI-assisted technologies in the writing process

Generative AI was used for English-language editing and to improve readability. The authors reviewed all AI-assisted text and take full responsibility for the content.

