## Supplementary material for "Auricular vagus nerve stimulation facilitates contextual fear extinction via insular–mPFC pathway without affecting reconsolidation": Sup

### 1 Supplementary information

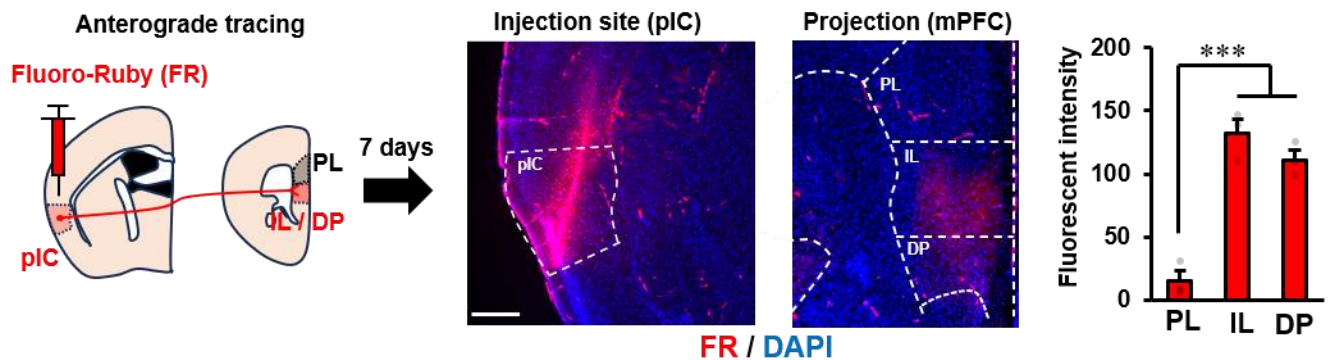

**Supplementary Fig. 1 The posterior insular cortex (pIC) preferentially projects to the** **infralimbic (IL) and dorsal peduncular (DP) subregions of the medial prefrontal cortex** **(mPFC). (a)** Experimental procedure. An anterograde tracer, Fluoro-Ruby (FR), was injected into the pIC, and FR-labelled axons in the mPFC were examined. **(b)** Representative image of the FR injection site in the pIC. **(c)** Representative image of FR-labelled axons in the mPFC subregions (prelimbic cortex [PL], IL, and DP). **(d)** Quantification of FR signal intensity in the mPFC subregions. One-way analysis of variance reveals a significant main effect of subregion ($F(2, 6) = 47.25, p < 0.001$). Tukey's post hoc test reveals a lower FR intensity in the PL than in the IL and DP ($***p < 0.001$), whereas the FR intensity does not differ between the IL and DP ($p$ $= 0.301$). Group sizes: $n = 3$.

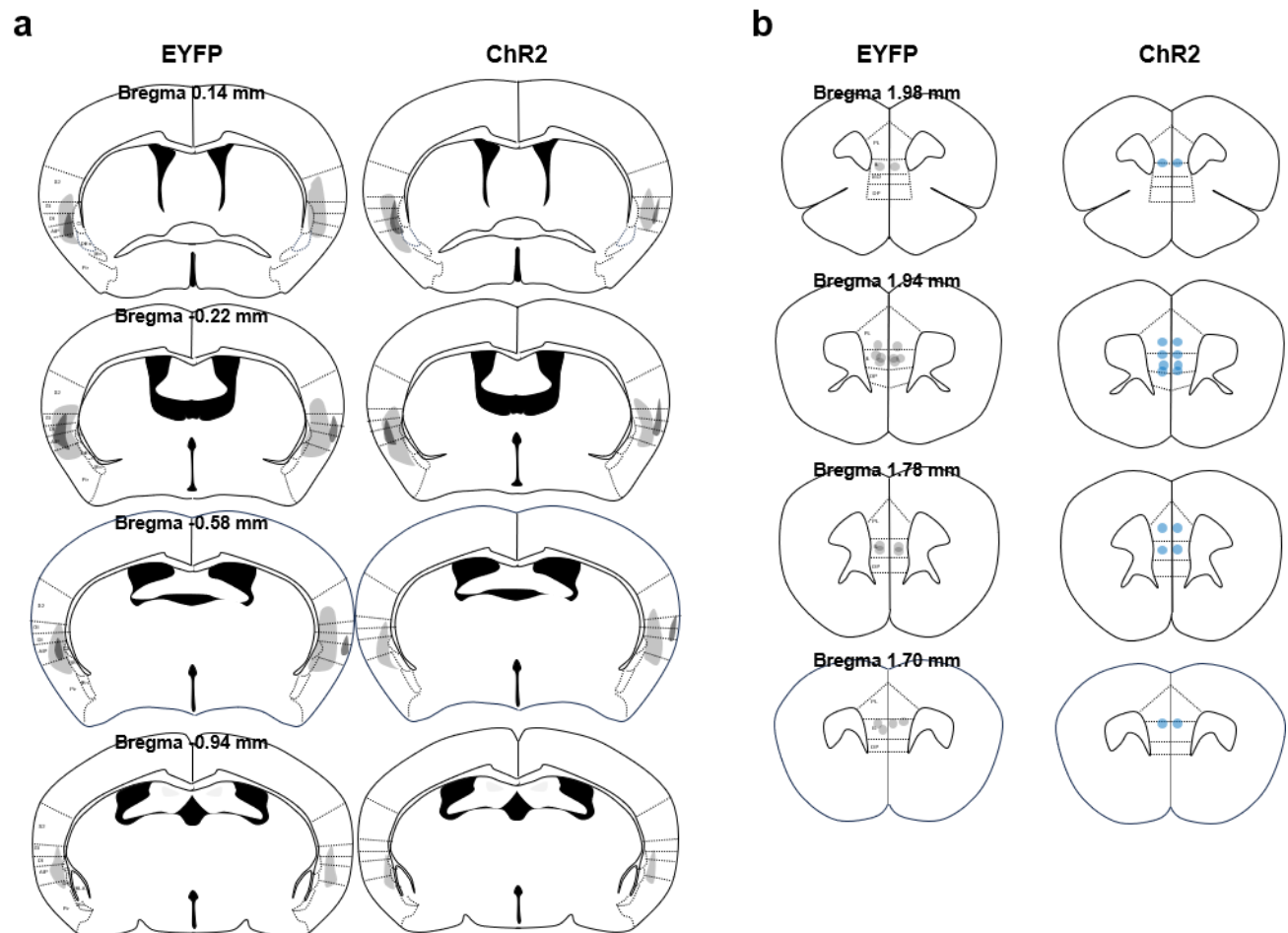

**Supplementary Fig. 2 Viral injection sites in the posterior insular cortex (pIC) and optic fibre placements in the infralimbic/dorsal peduncular cortex (IL/DP) for optogenetic activation experiments (related to Figure 6).** (a) Schematic maps show the minimum (light shading) and maximum (dark shading) spread of enhanced yellow fluorescent protein (EYFP) or channelrhodopsin-2 (ChR2)–EYFP expression in mice following adeno-associated virus injection into the pIC. (b) Locations of the optic fibres placed above the IL/DP in the mice for which data is shown in Figure 6. Each circle indicates an individual fibre placement, and colour intensity indicates the degree of overlap across animals.

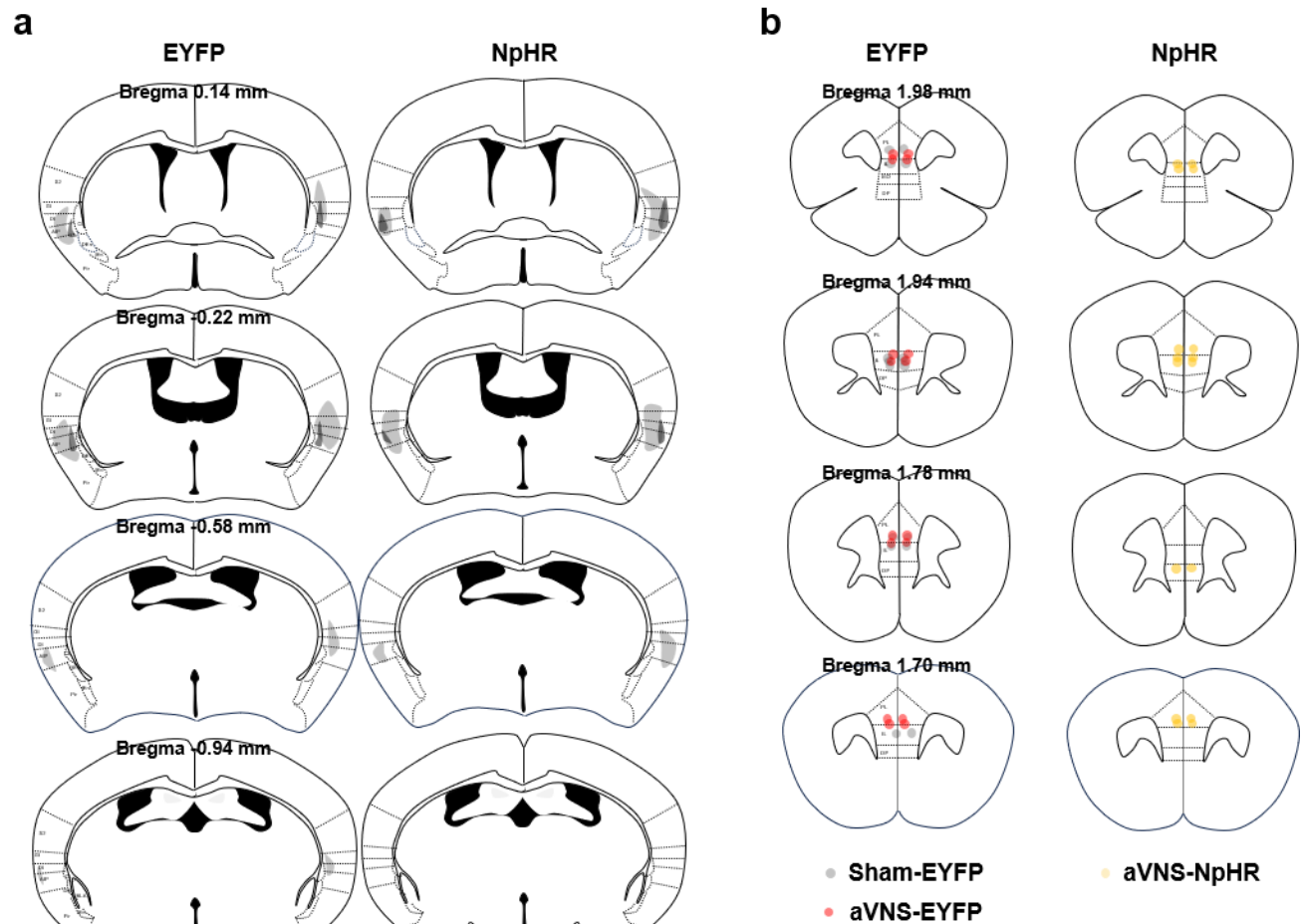

**Supplementary Fig. 3 Viral injection sites in the posterior insular cortex (pIC) and optic fibre placements in the infralimbic/dorsal peduncular cortex (IL/DP) for optogenetic inhibition experiments (related to Figure 7). (a) Schematic maps show the minimum (light shading) and maximum (dark shading) spread of EYFP or halorhodopsin (NpHR)–EYFP expression in mice following adeno-associated virus injection into the pIC. (b) Locations of the optic fibres placed above the IL/DP in the mice for which data is shown in Figure 7. Each circle indicates an individual fibre placement, and colour intensity indicates the degree of overlap across animals. aVNS, auricular vagus nerve stimulation.**

|  |  |  |  |  |
| --- | --- | --- | --- | --- |
| Fig. 2b | Repeated-measures ANOVA |  |  |  |
|  | Effects | F value | p value | Significance |
| | Treatment | $F(3, 26) = 0.423$ | $p = 0.988$ | n.s. |
| | Bin | $F(1, 26) = 367.330$ | $p < 0.001$ | *** |
| | Treatment × Bin | $F(3, 26) = 0.19$ | $p = 0.903$ | n.s. |
| Fig. 2c | Repeated-measures ANOVA |  |  |  |
|  | Effects | F value | p value | Significance |
| | Treatment | $F(3, 26) = 1.850$ | $p = 0.163$ | n.s. |
| | Bin | $F(2, 52) = 3.000$ | $p = 0.058$ | n.s. |
| | Treatment × Bin | $F(6, 52) = 3.070$ | $p = 0.012$ | * |
|  | Tukey's HSD post hoc test |  |  |  |
|  | Bin | Comparison | p value | Significance |
| | 1 min | aVNS-unstim vs. aVNS-stim | $p = 0.148$ | n.s. |
| | | Sham-stim vs. aVNS-stim | $p = 0.026$ | * |
| | | Sham-unstim vs. aVNS-stim | $p = 0.125$ | n.s. |
| | | Sham-stim vs. aVNS-unstim | $p = 0.962$ | n.s. |
| | | Sham-unstim vs. aVNS-unstim | $p = 0.997$ | n.s. |
| | | Sham-unstim vs. Sham-stim | $p = 0.995$ | n.s. |
| | 2 min | aVNS-unstim vs. aVNS-stim | $p = 0.145$ | n.s. |
| | | Sham-stim vs. aVNS-stim | $p = 0.069$ | n.s. |
| | | Sham-unstim vs. aVNS-stim | $p = 0.515$ | n.s. |
| | | Sham-stim vs. aVNS-unstim | $p = 0.999$ | n.s. |
| | | Sham-unstim vs. aVNS-unstim | $p = 0.926$ | n.s. |
| | | Sham-unstim vs. Sham-stim | $p = 0.869$ | n.s. |
| | 3 min | aVNS-unstim vs. aVNS-stim | $p = 0.977$ | n.s. |
| | | Sham-stim vs. aVNS-stim | $p = 0.790$ | n.s. |
| | | Sham-unstim vs. aVNS-stim | $p = 0.996$ | n.s. |
| | | Sham-stim vs. aVNS-unstim | $p = 0.629$ | n.s. |
| | | Sham-unstim vs. aVNS-unstim | $p = 0.949$ | n.s. |
| | | Sham-unstim vs. Sham-stim | $p = 0.945$ | n.s. |
| Fig. 2d | Repeated-measures ANOVA |  |  |  |
|  | Effects | F value | p value | Significance |
| | Treatment | $F(3, 26) = 0.120$ | $p = 0.946$ | n.s. |
| | Bin | $F(2, 52) = 3.650$ | $p = 0.033$ | * |
| | Treatment × Bin | $F(6, 52) = 1.110$ | $p = 0.368$ | n.s. |
| Fig. 2e | Repeated-measures ANOVA |  |  |  |
|  | Effects | F value | p value | Significance |
| | Treatment | $F(3, 36) = 0.040$ | $p = 0.988$ | n.s. |
| | Session | $F(3, 78) = 381.140$ | $p < 0.001$ | *** |
| | Treatment × Session | $F(9, 78) = 0.080$ | $p = 0.999$ | n.s. |

**Supplementary Table 1. Detailed statistical results for Figure 2.**

Repeated-measures analysis of variance (ANOVA) results for freezing behaviour shown in

Figure 2b–e, including main effects and interactions. Tukey's honestly significant difference post

33 hoc comparisons are reported for Figure 2c, where a significant Treatment × Bin interaction was
34 detected. Post hoc comparisons were performed only when a significant interaction or a
35 significant main effect was detected in the repeated-measures ANOVA. Significance is indicated
36 as n.s., not significant; \*$p < 0.05$; \*\*\*$p < 0.001$. aVNS, auricular vagus nerve stimulation.

37

|  |  |  |  |  |
| --- | --- | --- | --- | --- |
| Fig. 3b | Repeated-measures ANOVA |  |  |  |
|  | Effects | F value | p value | Significance |
|  | Treatment | F(3, 36) = 0.690 | p = 0.5668 | n.s. |
|  | Bin | F(1, 36) = 1169.80 | p < 0.001 | *** |
|  | Treatment × Bin | F(3, 36) = 1.190 | p = 0.3271 | n.s. |
| Fig. 3c | Repeated-measures ANOVA |  |  |  |
|  | Effects | F value | p value | Significance |
|  | Treatment | F(3, 36) = 11.72 | p < 0.001 | *** |
|  | Bin | F(5, 180) = 25.10 | p < 0.001 | *** |
|  | Treatment × Bin | F(15, 180) = 1.200 | p = 0.278 | n.s. |
|  | Tukey's HSD post hoc test |  |  |  |
|  | Bin | Comparison | p value | Significance |
|  | 1 min | aVNS-unstim vs. aVNS-stim | p = 0.009 | ** |
|  |  | Sham-stim vs. aVNS-stim | p = 0.002 | ** |
|  |  | Sham-unstim vs. aVNS-stim | p = 0.001 | ** |
|  |  | Sham-stim vs. aVNS-unstim | p = 0.927 | n.s. |
|  |  | Sham-unstim vs. aVNS-unstim | p = 0.835 | n.s. |
|  | 2 min | Sham-unstim vs. Sham-stim | p = 0.996 | n.s. |
|  |  | aVNS-unstim vs. aVNS-stim | p < 0.001 | *** |
|  |  | Sham-stim vs. aVNS-stim | p < 0.001 | *** |
|  |  | Sham-unstim vs. aVNS-stim | p = 0.001 | ** |
|  |  | Sham-stim vs. aVNS-unstim | p = 0.974 | n.s. |
|  | 3 min | Sham-unstim vs. aVNS-unstim | p = 0.999 | n.s. |
|  |  | Sham-unstim vs. Sham-stim | p = 0.938 | n.s. |
|  |  | aVNS-unstim vs. aVNS-stim | p = 0.001 | ** |
|  |  | Sham-stim vs aVNS-stim | p < 0.001 | *** |
|  |  | Sham-unstim vs aVNS-stim | p = 0.021 | * |
|  | 4 min | Sham-stim vs aVNS-unstim | p = 0.695 | n.s. |
|  |  | Sham-unstim vs aVNS-unstim | p = 0.721 | n.s. |
|  |  | Sham-unstim vs Sham-stim | p = 0.159 | n.s. |
|  |  | aVNS-unstim vs aVNS-stim | p = 0.042 | * |
|  |  | Sham-stim vs aVNS-stim | p < 0.001 | *** |
|  | 5 min | Sham-stim vs aVNS-stim | p = 0.300 | n.s. |
|  |  | Sham-unstim vs aVNS-unstim | p = 0.925 | n.s. |
|  |  | Sham-unstim vs Sham-stim | p = 0.656 | n.s. |
|  |  | aVNS-unstim vs aVNS-stim | p = 0.004 | * |
|  |  | Sham-stim vs aVNS-stim | p = 0.009 | ** |
|  | 6 min | Sham-unstim vs aVNS-stim | p = 0.004 | ** |
|  |  | Sham-stim vs aVNS-stim | p = 0.003 | ** |
|  |  | Sham-unstim vs aVNS-stim | p = 0.019 | * |
|  |  | Sham-stim vs aVNS-unstim | p = 1.000 | n.s. |
|  |  | Sham-unstim vs aVNS-unstim | p = 0.921 | n.s. |
|  |  | Sham-unstim vs Sham-stim | p = 0.887 | n.s. |

|  |  |  |  |  |
| --- | --- | --- | --- | --- |
| Fig. 3d | Repeated-measures ANOVA |  |  |  |
|  | Effects | F value | p value | Significance |
|  | Treatment | F(3, 36) = 7.540 | p < 0.001 | *** |
|  | Bin | F(2, 72) = 4.690 | p = 0.012 | * |
|  | Treatment × Bin | F(6, 72) = 0.810 | p = 0.566 | n.s. |
|  | Tukey's HSD post hoc test |  |  |  |
|  | Bin | Comparison | p value | Significance |
|  | 1 min | aVNS-unstim vs. aVNS-stim | p = 0.002 | ** |
|  |  | Sham-stim vs. aVNS-stim | p = 0.015 | * |
|  |  | Sham-unstim vs. aVNS-stim | p = 0.009 | ** |
|  |  | Sham-stim vs. aVNS-unstim | p = 0.876 | n.s. |
|  |  | Sham-unstim vs. aVNS-unstim | p = 0.946 | n.s. |
|  | 2 min | Sham-unstim vs. Sham-stim | p = 0.997 | n.s. |
|  |  | aVNS-unstim vs. aVNS-stim | p = 0.016 | * |
|  |  | Sham-stim vs. aVNS-stim | p = 0.049 | * |
|  |  | Sham-unstim vs. aVNS-stim | p = 0.020 | * |
|  |  | Sham-stim vs. aVNS-unstim | p = 0.965 | n.s. |
|  | 3 min | Sham-unstim vs. aVNS-unstim | p = 1.000 | n.s. |
|  |  | Sham-unstim vs. Sham-stim | p = 0.980 | n.s. |
|  |  | aVNS-unstim vs aVNS-stim | p = 0.008 | ** |
|  |  | Sham-stim vs aVNS-stim | p = 0.003 | ** |
|  |  | Sham-unstim vs aVNS-stim | p = 0.009 | ** |
| Fig. 3e | Repeated-measures ANOVA |  |  |  |
|  | Effects | F value | p value | Significance |
|  | Treatment | F(3, 36) = 6.52 | p = 0.001 | ** |
|  | Session | F(3, 108) = 546.69 | p < 0.001 | *** |
|  | Treatment × Session | F(9, 108) = 6.61 | p < 0.001 | *** |
|  | Tukey's HSD post hoc test |  |  |  |
|  | Session | Comparison | p value | Significance |
|  | Day1 preshock | aVNS-unstim vs. aVNS-stim | p = 0.999 | n.s. |
|  |  | Sham-stim vs. aVNS-stim | p = 1.000 | n.s. |
|  |  | Sham-unstim vs. aVNS-stim | p = 0.981 | n.s. |
|  |  | Sham-stim vs. aVNS-unstim | p = 1.000 | n.s. |
|  |  | Sham-unstim vs. aVNS-unstim | p = 0.917 | n.s. |
|  | Day1 postshock | Sham-unstim vs. Sham-stim | p = 0.976 | n.s. |
|  |  | aVNS-unstim vs. aVNS-stim | p = 0.539 | n.s. |
|  |  | Sham-stim vs. aVNS-stim | p = 0.841 | n.s. |
|  |  | Sham-unstim vs. aVNS-stim | p = 1.000 | n.s. |
|  |  | Sham-stim vs. aVNS-unstim | p = 0.954 | n.s. |
|  | Day2 | Sham-unstim vs. aVNS-unstim | p = 0.493 | n.s. |
|  |  | Sham-unstim vs. Sham-stim | p = 0.804 | n.s. |
|  |  | aVNS-unstim vs aVNS-stim | p < 0.001 | *** |
|  |  | Sham-stim vs aVNS-stim | p < 0.001 | *** |
|  |  | Sham-unstim vs aVNS-stim | p < 0.001 | *** |
|  | Day3 | Sham-stim vs aVNS-unstim | p = 0.819 | n.s. |
|  |  | Sham-unstim vs aVNS-unstim | p = 1.000 | n.s. |
|  |  | Sham-unstim vs Sham-stim | p = 0.834 | n.s. |
|  |  | aVNS-unstim vs aVNS-stim | p = 0.003 | ** |
|  |  | Sham-stim vs aVNS-stim | p = 0.015 | * |
|  |  | Sham-unstim vs aVNS-stim | p = 0.008 | ** |
|  |  | Sham-stim vs aVNS-unstim | p = 0.930 | n.s. |
|  |  | Sham-unstim vs aVNS-unstim | p = 0.930 | n.s. |
|  |  | Sham-unstim vs Sham-stim | p = 0.995 | n.s. |

### Supplementary Table 2. Detailed statistical results for Figure 3.

Repeated-measures analysis of variance (ANOVA) results for freezing behaviour shown in Figure 3b–e, including main effects and interactions. Tukey's honestly significant difference post hoc comparisons are reported for panels in which a significant interaction or a significant main effect was detected. Post hoc comparisons were performed only when a significant interaction or a significant main effect was detected in the repeated-measures ANOVA. Significance is indicated as n.s., not significant; \*p < 0.05; \*\*p < 0.01; \*\*\*p < 0.001. aVNS, auricular vagus nerve stimulation.

|  |  |  |  |  |
| --- | --- | --- | --- | --- |
| Fig. 4b | Repeated-measures ANOVA |  |  |  |
|  | Effects | F value | p value | Significance |
|  | Treatment | F(1, 16) = 0.018 | p = 0.894 | n.s. |
|  | Bin | F(1, 16) = 792.607 | p < 0.001 | *** |
| Fig. 4c | Repeated-measures ANOVA |  |  |  |
|  | Effects | F value | p value | Significance |
|  | Treatment | F(1, 16) = 2.538 | p = 0.1307 | n.s. |
|  | Bin | F(5, 80) = 44.657 | p < 0.001 | *** |
|  | Treatment × Bin | F(5, 80) = 8.790 | p < 0.001 | *** |
|  | post hoc t-test |  |  |  |
|  | Bin | t value | p value | Significance |
|  | 1 min | t(16) = -0.014 | p = 0.989 | n.s. |
|  | 2 min | t(16) = 0.759 | p = 0.459 | n.s. |
|  | 3 min | t(16) = 0.688 | p = 0.501 | n.s. |
| Fig. 4d | Repeated-measures ANOVA |  |  |  |
|  | Effects | F value | p value | Significance |
|  | Treatment | F(1, 16) = 25.661 | p < 0.001 | *** |
|  | Bin | F(2, 32) = 5.499 | p = 0.009 | * |
|  | Treatment × Bin | F(2, 32) = 0.149 | p = 0.862 | n.s. |
|  | post hoc t-test |  |  |  |
|  | Bin | t value | p value | Significance |
|  | 1 min | t(16) = -3.360 | p = 0.004 | ** |
|  | 2 min | t(16) = -5.470 | p < 0.001 | *** |
|  | 3 min | t(16) = -4.470 | p < 0.001 | *** |
| Fig. 4e | Repeated-measures ANOVA |  |  |  |
|  | Effects | F value | p value | Significance |
|  | Treatment | F(1, 16) = 5.517 | p = 0.032 | * |
|  | Session | F(4, 64) = 204.330 | p < 0.001 | *** |
|  | Treatment × Session | F(4, 64) = 7.280 | p < 0.001 | *** |
|  | post hoc t-test |  |  |  |
|  | Session | t value | p value | Significance |
|  | Day1 preshock | t(16) = 0.219 | p = 0.829 | n.s. |
|  | Day1 postshock | t(16) = 0.099 | p = 0.923 | n.s. |
|  | Day2 Early | t(16) = 0.510 | p = 0.617 | n.s. |

|  |  |  |  |  |
| --- | --- | --- | --- | --- |
| Fig. 4g | Repeated-measures ANOVA |  |  |  |
|  | Effects | F value | p value | Significance |
|  | Treatment | F(1, 18) = 0.013 | p = 0.912 | n.s. |
|  | Bin | F(1, 18) = 1165.480 | p < 0.001 | *** |
| Fig. 4h | Repeated-measures ANOVA |  |  |  |
|  | Effects | F value | p value | Significance |
|  | Treatment | F(1, 18) = 3.279 | p = 0.0869 | n.s. |
|  | Bin | F(5, 90) = 26.288 | p < 0.001 | *** |
|  | Treatment × Bin | F(5, 90) = 3.358 | p < 0.008 | ** |
|  | post hoc t-test |  |  |  |
|  | Bin | t value | p value | Significance |
|  | 1 min | t(18) = -3.970 | p < 0.001 | *** |
|  | 2 min | t(18) = -2.120 | p = 0.048 | * |
|  | 3 min | t(18) = -2.650 | p = 0.016 | * |
| Fig. 4i | Repeated-measures ANOVA |  |  |  |
|  | Effects | F value | p value | Significance |
|  | Treatment | F(1, 18) = 0.004 | p = 0.953 | n.s. |
|  | Bin | F(2, 36) = 4.451 | p = 0.019 | * |
|  | Treatment × Bin | F(2, 36) = 0.276 | p = 0.760 | n.s. |
|  | post hoc t-test |  |  |  |
|  | Bin | t value | p value | Significance |
|  | 1 min | t(18) = -0.071 | p = 0.944 | n.s. |
|  | 2 min | t(18) = 0.281 | p = 0.782 | n.s. |
|  | 3 min | t(18) = -0.362 | p = 0.722 | n.s. |
| Fig. 4j | Repeated-measures ANOVA |  |  |  |
|  | Effects | F value | p value | Significance |
|  | Treatment | F(1, 18) = 1.470 | p = 0.241 | n.s. |
|  | Session | F(4, 72) = 213.660 | p < 0.001 | *** |
| Fig. 4k | Repeated-measures ANOVA |  |  |  |
|  | Treatment × Session | F(4, 72) = 2.318 | p = 0.065 | n.s. |

#### Supplementary Table 3. Detailed statistical results for Figure 4.

Repeated-measures analysis of variance (ANOVA) results for freezing behaviour shown in Figure 4b–e and g–j, including main effects and interactions. Post hoc comparisons are reported for panels in which a significant interaction or a significant main effect was detected. Post hoc tests were performed only when the repeated-measures ANOVA indicated a significant interaction or a significant main effect. Significance is indicated as n.s., not significant; \*$p < 0.05$; \*\*$p < 0.01$; \*\*\*$p < 0.001$. aVNS, auricular vagus nerve stimulation.

|  |  |  |  |  |
| --- | --- | --- | --- | --- |
| Fig. 6d | Repeated-measures ANOVA |  |  |  |
|  | Effects | F value | p value | Significance |
| | Treatment | $F(1, 15) = 0.009$ | $p = 0.927$ | n.s. |
| | Bin | $F(1, 15) = 314.305$ | $p < 0.001$ | *** |
| | Treatment × Bin | $F(1, 15) = 0.011$ | $p = 0.912$ | n.s. |
| Fig. 6e | Repeated-measures ANOVA |  |  |  |
|  | Effects | F value | p value | Significance |
| | Treatment | $F(1, 15) = 4.800$ | $p = 0.0447$ | * |
| | Bin | $F(5, 75) = 88.772$ | $p < 0.001$ | *** |
| | Treatment × Bin | $F(5, 75) = 20.862$ | $p < 0.001$ | *** |
|  | post hoc t-test |  |  |  |
|  | Bin | t value | p value | Significance |
| | 1 min | $t(15) = -0.307$ | $p = 0.763$ | n.s. |
| | 2 min | $t(15) = -0.354$ | $p = 0.728$ | n.s. |
| | 3 min | $t(15) = 0.409$ | $p = 0.688$ | n.s. |
| | 4 min | $t(15) = -3.283$ | $p = 0.005$ | ** |
| | 5 min | $t(15) = -4.325$ | $p < 0.001$ | *** |
| | 6 min | $t(15) = -3.536$ | $p = 0.003$ | ** |
| Fig. 6f | Repeated-measures ANOVA |  |  |  |
|  | Effects | F value | p value | Significance |
| | Treatment | $F(1, 15) = 14.396$ | $p = 0.002$ | ** |
| | Bin | $F(2, 30) = 11.675$ | $p < 0.001$ | *** |
| | Treatment × Bin | $F(2, 30) = 1.313$ | $p = 0.284$ | n.s. |
|  | post hoc t-test |  |  |  |
|  | Bin | t value | p value | Significance |
| | 1 min | $t(15) = -3.762$ | $p = 0.002$ | ** |
| | 2 min | $t(15) = -3.422$ | $p = 0.004$ | ** |
| | 3 min | $t(15) = -3.773$ | $p = 0.002$ | ** |
| Fig. 6g | Repeated-measures ANOVA |  |  |  |
|  | Effects | F value | p value | Significance |
| | Treatment | $F(1, 15) = 4.507$ | $p = 0.0508$ | n.s. |
| | Session | $F(4, 60) = 187.846$ | $p < 0.001$ | *** |
| | Treatment × Session | $F(4, 60) = 15.415$ | $p < 0.001$ | *** |
|  | post hoc t-test |  |  |  |
|  | Session | t value | p value | Significance |
| | Day1 preshock | $t(15) = -0.405$ | $p = 0.692$ | n.s. |
| | Day1 postshock | $t(15) = 0.098$ | $p = 0.923$ | n.s. |
| | Day2 Early | $t(15) = -0.101$ | $p = 0.921$ | n.s. |
| | Day2 Late | $t(15) = -3.883$ | $p = 0.001$ | ** |
| | Day3 Total | $t(15) = -3.794$ | $p = 0.002$ | ** |

**Supplementary Table 4. Detailed statistical results for Figure 6.**

58 Repeated-measures analysis of variance (ANOVA) results for freezing behaviour shown in
59 Figure 6d–g, including main effects and interactions. Post hoc comparisons (unpaired t-tests) are
60 reported for panels in which a significant interaction or a significant main effect was detected.
61 Post hoc tests were performed only when the repeated-measures ANOVA indicated a significant
62 interaction or a significant main effect. Significance is indicated as n.s., not significant; \*p <
63 0.05; \*\*p < 0.01; \*\*\*p < 0.001.

64

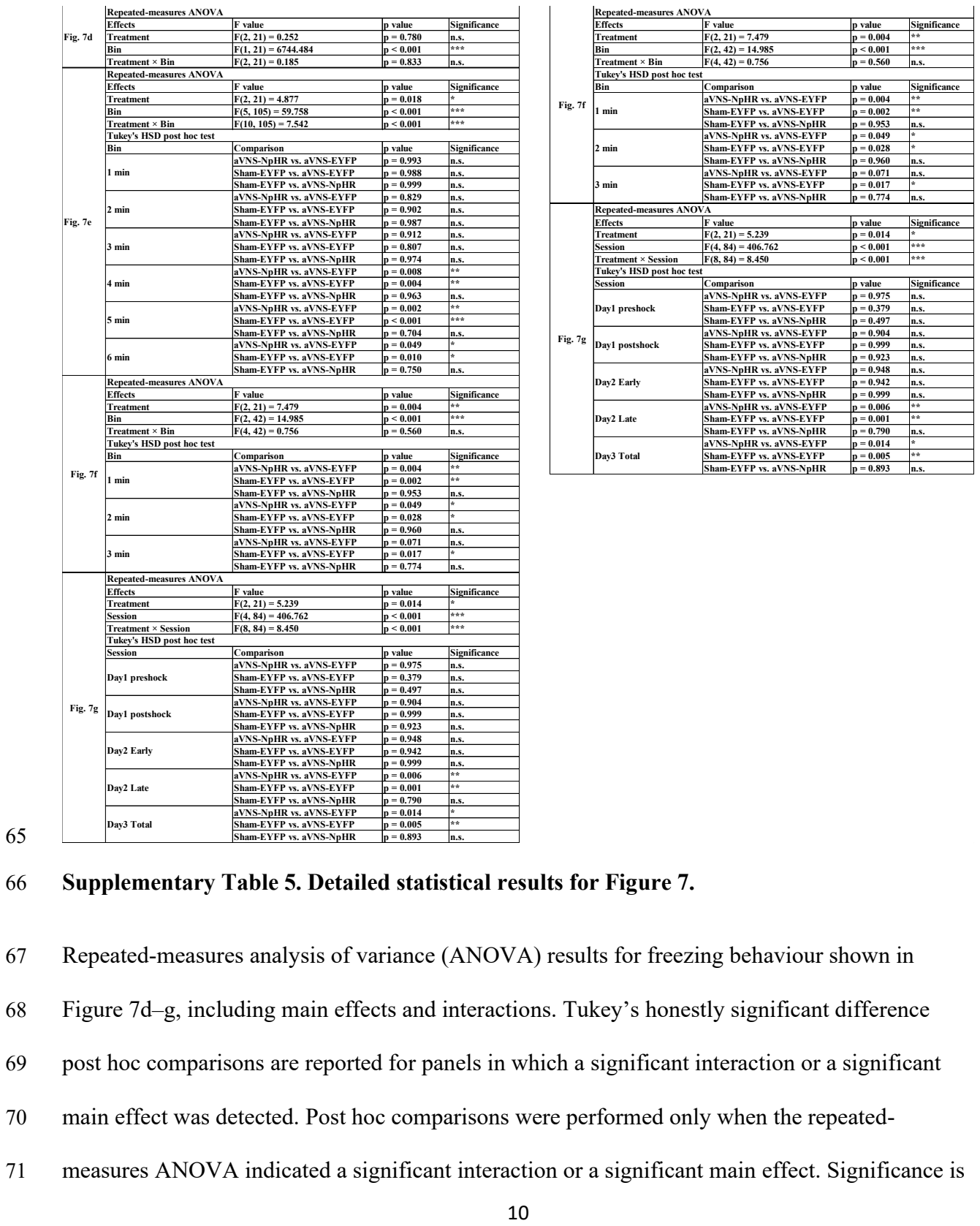

72 indicated as n.s., not significant; \*$p < 0.05$; \*\*$p < 0.01$; \*\*\*$p < 0.001$. aVNS, auricular vagus
73 nerve stimulation; EYFP, enhanced yellow fluorescent protein; NpHR, halorhodopsin.

74
